# Structural and coding variation in *PHYTOCHROMES A* and *C* underlies differences in flowering time and shade avoidance in wheat

**DOI:** 10.64898/2026.04.28.721246

**Authors:** Sai Thejas Babanna, Yogev Burko, Björn Christopher Willige, Gizaw Wolde, Thorsten Schnurbusch, Guy Golan

**Affiliations:** Leibniz Institute of Plant Genetics and Crop Plant Research (IPK), Corrensstr. 3, OT Gatersleben, 06466 Seeland, Germany; The Institute of Plant Sciences, Agricultural Research Organization, Volcani Center, Rishon LeZion 7505101, Israel; Department of Soil and Crop Sciences, College of Agricultural Sciences, Colorado State University, Fort Collins, CO 80521, USA; The Robert H. Smith Faculty of Agriculture, Food & Environment, The Hebrew University of Jerusalem, Rehovot 7610001, Israel; Faculty of Natural Sciences III, Institute of Agricultural and Nutritional Sciences, Martin Luther University Halle-Wittenberg, 06120 Halle, Germany; Cluster of Excellence on Plant Sciences (CEPLAS) ‘SMART Plants for Tomorrow’s Needs’

**Keywords:** Shade avoidance, phytochrome A, phytochrome C, wheat, flowering time

## Abstract

Plant architecture and developmental timing are influenced by light availability, especially in high-density cropping systems, where canopy shading modifies both light intensity and spectral quality. Despite their ecological and agronomical importance, the genetic basis of these responses in wheat remains poorly understood. Here, using a Recombinant Inbred Lines (RILs) population, we investigated phenological and morphological traits under sunlight and simulated canopy shade. We identify a major QTL on chromosome 5A with light-dependent allelic effects, indicating genotype-by-environment variation in developmental responses. This QTL corresponds to a structural rearrangement, consistent with an inversion in the wild emmer reference genome encompassing *PHYTOCHROME C* (*PHYC-A*) and *VERNALIZATION-1* (*VRN-A1*), as well as coding polymorphism in *PHYC-A*. Analysis of a tetraploid wheat diversity panel further showed that natural variation at *PHYC-A* and an early stop codon in the *BB* genome copy of *PHYTOCHROME A* (*PHYA-B*) on chromosome 4B are associated with differences in heading time. Functional analysis using TILLING-derived phytochrome mutants confirms distinct and complementary roles for *PHYA* and *PHYC* in regulating flowering time, plant height, and leaf elongation under simulated canopy shade. These findings highlight the contribution of phytochrome variation to developmental plasticity under canopy-like light environment, thereby extending model insights to agronomically relevant conditions.

## Introduction

Cereal crops are typically cultivated in high-density stands to promote rapid canopy closure, which enhances weed suppression, improves light interception, and maximizes dry matter accumulation per unit area (Mhlanga *et al*., 2016; Zhang *et al*., 2021, 2023). However, while high-density planting is common, it also imposes growth limitations on individual plants due to intense competition for resources (Weiner and Freckleton, 2010). Comparative analyses across species have shown that plant responses to high density resemble responses to low light, indicating that light competition plays a role in shaping plant physiology, morphology, anatomy, growth, and reproduction (Poorter *et al*., 2016, 2019; Postma *et al*., 2021).

The extent to which plant traits respond to light availability or planting density varies both among taxa and within species (Poorter *et al*., 2019; Postma *et al*., 2021). This variation has important implications for fitness across different ecological scenarios (Ballaré and Pierik, 2017). Shade avoidance responses are also prevalent in agricultural settings, where traits such as reduced tillering, stem elongation, and altered leaf morphology can have either beneficial or detrimental effects on productivity, depending on the specific crop and cropping system (Ballaré *et al*., 1991; Ballaré and Casal, 2000; López Pereira *et al*., 2017). However, the extent to which natural variation in shade responses shapes plant development and architecture in realistic field conditions, and the genetic mechanisms driving shade-induced developmental plasticity, remain largely unknown.

Plants sense light using various photoreceptors, among which phytochromes play a central role in sensing canopy-associated light cues. Phytochromes respond to changes in the red to far-red ratio that arise from selective absorbance of different phytochrome forms and the light reflected within plant canopies (Batschauer, 1998; Briggs and Olney, 2001; Quail, 2002). In grasses, the phytochrome gene family contains three members *PHYA*, *PHYB*, and *PHYC*, with PHYB and PHYC proteins regulating flowering time in wheat (*Triticum* sp.), barley (*Hordeum vulgare* L.) and maize (*Zea mays* L.) (Nishida *et al*., 2013; Chen *et al*., 2014; Li *et al*., 2020; Alvarez *et al*., 2023). In wheat plants, the role of phytochromes in regulating flowering time is complex and involves both photoperiodic and vernalization pathways. Photoperiodic flowering is regulated by the interaction between phytochromes and other genes, such as the *PHOTOPERIOD1* (*PPD1*), *VERNALIZATION-1* (*VRN-1*), *FLOWERING LOCUS T-1* (*FT-1*) and *EARLY FLOWERING 3* (*ELF3*), which initiate the flowering process in wheat (Chen *et al*., 2014; Alvarez *et al*., 2023).

Most studies of shade responses have been conducted in model species, often focusing on the spectral composition of low-intensity light under controlled growth conditions. While these approaches have been invaluable for dissecting light signaling pathways, they typically decouple light cues from the developmental, physiological, and phenological context in which shade is perceived in crop fields. Experimental systems that bridge this gap, capturing key features of the canopy environment while retaining sufficient environmental control remain relatively rare. Simulated canopy shade treatments (Golan *et al*., 2023) provide an intermediate level of realism, allowing shade-mediated plasticity to be examined under conditions that better reflect crop canopies.

In this study, we investigate the genetic basis of shade responsiveness for tetraploid wheat morphology and phenology in a simulated canopy shade setup. We identify previously uncharacterized phytochrome polymorphisms associated with developmental and morphological responses to shade, and we provide functional support for the role of phytochromes using mutant lines. Together, this study enables us to assess how variation in light signaling pathways contributes to phenotypic variation under controlled yet more canopy- and agronomically relevant conditions.

## Materials and Methods

### Plant materials and growth conditions

Recombinant Inbred Lines (RILs) were derived from a cross between the durum cultivar (*Triticum turgidum* ssp. *durum*) Svevo (Maccaferri *et al*., 2019) and the wild emmer (*Triticum turgidum* ssp. *dicoccoides*) accession Zavitan (Avni *et al*., 2017) that differ in plant architecture, phenology and adaptation to cultivation. Genotyping of the RILs was carried out with the Illumina iSelect 90K SNP array (Avni *et al*., 2014; Wang *et al*., 2014) and the genetic map was constructed as previously described (Nave *et al*., 2016; Golan *et al*., 2024).

Tilling mutants were derived from the tetraploid wheat *Triticum turgidum* ssp. durum cultivar Kronos mutant population (Krasileva *et al*., 2017). Four phytochrome knockout mutants were selected to investigate the roles of *PHYA* and *PHYC* genes, and were backcrossed two to three times to Kronos to generate Near-Isogenic Lines (NILs). The *phyA-A* (Kronos2134-derived) and *phyA-B* (Kronos499-derived) NILs contain early stop codons, resulting in truncated proteins of 522 and 458 amino acids, respectively, both of which lack the PHY, PAS-related, and histidine kinase-related domains. Functional analysis of *PHYC* was performed using two NIL mutants: *phyC-A* (Kronos2327-derived), which harbors a mutation in the donor splice site at the end of exon 2, leading to a premature stop codon and loss of the C-terminal 179 amino acids; and *phyC-B* (Kronos807-derived), which carries an amino acid substitution in the GAF domain that disrupts protein function (Chen *et al*., 2014).

RILs were grown in 2021 in two adjacent glasshouses under natural soil and environmental conditions, following the protocol described previously (Golan *et al*., 2024). Each glasshouse represented a distinct light environment (natural sunlight or simulated canopy shade), and individual plants constituted the experimental units for phenotypic analyses. Seedlings were germinated under controlled conditions (16 h light / 8 h dark, 20/16°C) and transplanted at the two-leaf stage into glasshouses arranged in an alpha-lattice design. A simulated canopy-shade treatment was imposed using a green plastic filter (Lee122 Fern Green), which reduced blue and red light by ∼80%, lowered the red/far-red ratio from 1.13 to 0.28, and decreased PAR by ∼55%. Plants were spaced to allow single-plant phenotyping, were irrigated regularly, and fertilized during stem elongation. Additional details on soil composition, glasshouse structure, and management practices are provided in Golan et al., (2024). This approach allowed us to apply canopy shade while maintaining suitable plant spacing, thereby diminishing the confounding influences of belowground competition. Tilling-derived near-isogenic lines were cultivated in the same glasshouses under similar conditions in the following year (2022).

### Phenotyping

All the phenotypic data were recorded from the main culm to enable consistent comparisons among genotypes. Traits were selected to capture both early and late vegetative responses and integrated developmental outcomes. The number of days to heading was recorded when the spike emerged half out of the leaf sheaths. Plant height, specific stem length (stem length/stem dry mass), flag leaf size, and final leaf number were measured at maturity, while the size of Leaf 2 and Leaf 3 and their spacing were recorded at the 4-leaf stage, which was marked on the main culm with a plastic straw. Tillers were counted during the early booting stage.

A mixed linear model was applied to evaluate the main effects of light environment and genotype, their interaction, and the spatial block effect on the phenotypic traits of the RIL population using the JMP 16 software (SAS Institute) as:

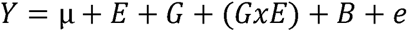

where the intercept (μ), and the light environment (E) are fixed effects, and the RIL (G), G x E interactions and the block as random effects.

### Tissue collection, RNA extraction and RT-qPCR

Expression analysis was designed to assess whether natural variation in phytochromes is associated with shifts in the temporal expression patterns of key flowering-time genes. Three groups of recombinant inbred lines (RILs) were selected for reverse transcription quantitative PCR (RT-qPCR) analysis based on the allelic combinations of flowering time QTL at four loci including the *PHYA-B* and *PHYC-A* loci, as explained in the Results section. Four to five independent RILs representing each allelic group were used, with each RIL considered a single biological replicate. To assess rhythmic gene expression in young leaves, the fourth leaf was harvested at five time points over a 24-hour period: 6, 10, 18, 22, and 2 hours (Zeitgeber Time). The fourth leaf was selected as a developmentally comparable tissue suitable for assessing light-responsive gene expression prior to floral transition. For RNA sampling, plants were grown in controlled growth chambers under the same photoperiod, temperature, and soil conditions as described above, with a light intensity of 270 µE m□² s□¹ and 70% relative humidity. Seeds from each RIL were sown in 96-well trays containing the same soil compost mix and grown in these trays until tissue collection for RNA extraction was completed at four-leaf stage. Total RNA was extracted using TRIzol reagent (Invitrogen, US) and precipitated with isopropanol. Residual genomic DNA was removed by treatment with DNase I (New England Biolabs, US). First-strand complementary DNA (cDNA) was synthesized from total RNA using the SuperScript III Reverse Transcriptase Kit (Invitrogen, US). Quantitative real-time PCR was performed using SYBR Green Master Mix (Thermo Fisher Scientific, US) on the ABI Prism 7900HT Sequence Detection System (Applied Biosystems, US). For each gene, three technical replicates were run per biological replicate. The 2-^ΔΔCT^ method (Livak and Schmittgen, 2001) was used to normalize and calibrate transcript values relative to *TaActin*. Primer sequences used for RT-qPCR are listed in Table S1.

### Genetic analysis

Best Linear Unbiased Estimator (BLUE) calculation and QTL analysis were conducted as described in Golan et al., (2024). Briefly, prior to QTL analysis, phenotypic values for each RIL within each light environment were adjusted using a two-stage approach implemented in Genstat (VSN International) to obtain unbiased genotype estimates within each environment while preserving appropriate error structures for subsequent QTL mapping. In the first stage, replication and block were fitted as random terms to estimate within-environment variance components. In the second stage, genotypes were fitted as fixed effects to obtain BLUEs for subsequent QTL mapping. QTL analysis followed the multi-environment trial framework described in Malosetti et al., 2013. The optimal variance–covariance structure for genotype effects within and between environments was selected using the Schwarz Information Criterion (SIC). Genome-wide scans were performed using simple interval mapping (SIM) with a maximum step size of 10 cM. Significant loci were incorporated as cofactors in composite interval mapping (CIM) until QTL profiles stabilized. QTL significance thresholds (α = 0.05) were determined according to the number of effectively independent tests (Li and Ji, 2005). Final multi-QTL models decomposed each locus into its main additive effect and the QTL × environment (light treatment) interaction:

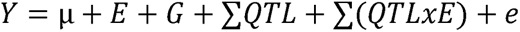

Where E is the fixed light-environment effect, G is the random RIL effect, ∑QTL is the total additive effect of each QTL, ∑(QTLxE) represents the total QTL effects x environment interactions, and e is the residual term structured according to the selected variance–covariance model.

### Genotypic profiling of *PHYC-A*, *PHYA-B* and *VRN-A1* in a tetraploid diversity panel

The tetraploid diversity panel was grown under controlled long day (16:8) conditions, as described by (Wolde *et al*., 2021). Heading time was recorded when half of the spike emerged from the flag leaf sheath. Sequence polymorphisms identified in *PHYC-A*, *PHYA-B*, and *VRN-A1* between Svevo and Zavitan were used to design PCR-based markers, which were subsequently applied to genotype 339 tetraploid wheat accessions including wild (*T. turgidum* ssp. *dicoccoides*) and domesticated emmer (*T. turgidum* ssp. *dicoccum*), miracle wheat (*T. turgidum* ssp. *compositum*), and durum wheat (*T. turgidum* ssp. *durum*) for the corresponding alleles. Details of the primers and PCR are provided in Table S1.

## Results

### Shade induces variation in plant architecture and phenology

To assess how wheat responds to canopy shade, we examined phenological and architectural traits in a RIL population grown under natural sunlight and simulated canopy shade. The RILs were assumed to differ in their shade response, due to the different selection forces acting on wild emmer plants in their natural habitat vs the strong artificial selection for productivity in managed environments during crop breeding. Traits were selected to capture both early vegetative responses and integrated developmental outcomes. Specifically, we examined a set of relevant traits, including plant height, specific stem length (stem length divided by weight), tiller number, leaf length, leaf spacing during early growth (the vertical distance between the second and third leaves), and flag leaf width. We also quantified the final leaf number on the main culm and heading time.

We used mixed linear models to examine shading effects, the genotype effect, and their interaction. Shading significantly increased the length of the early developing leaves and their vertical spacing (P<0.0001), as well as the width of the flag leaf (Table S2). Shading slightly reduced the final plant height but increased specific stem length, suggesting that competition for light under low light availability induces slenderness, which helps wheat plants reach the light. Shading also accelerated heading time and reduced leaf and tiller numbers. As expected, the genotype accounted for most of the phenotypic variation explained in the examined traits. However, the genotype-by-light environment interaction was significant for heading time, leaf spacing, and tiller number. These results suggest that in the studied population, genetic variation in phenotypic plasticity to shade mainly involves variation in developmental timing (Table S2).

### Identification of shade-induced plasticity loci

Correlation analysis revealed strong associations among the studied traits (Fig. S1). We therefore used Principal Component (PC) analysis to simplify interpretation and highlight major patterns in trait expression across the light environments. PC1 accounted for ∼34% of the phenotypic variation and was mainly loaded with tiller number, leaf number, days to heading, and leaf spacing and was therefore interpreted as a composite developmental axis (Fig. S2). The separation of shaded and sunlight phenotypes along PC1 (Fig. 1a) suggests that this component captures trait variation associated with the shade response. PC2 captured variation in leaf size and was significantly influenced by the light environment, with shading promoting leaf enlargement. PC3 showed differences between the light environments and was mainly loaded with flag leaf length, specific stem length, and final plant height (Figs. 1a, S3).

**Fig. 1.**
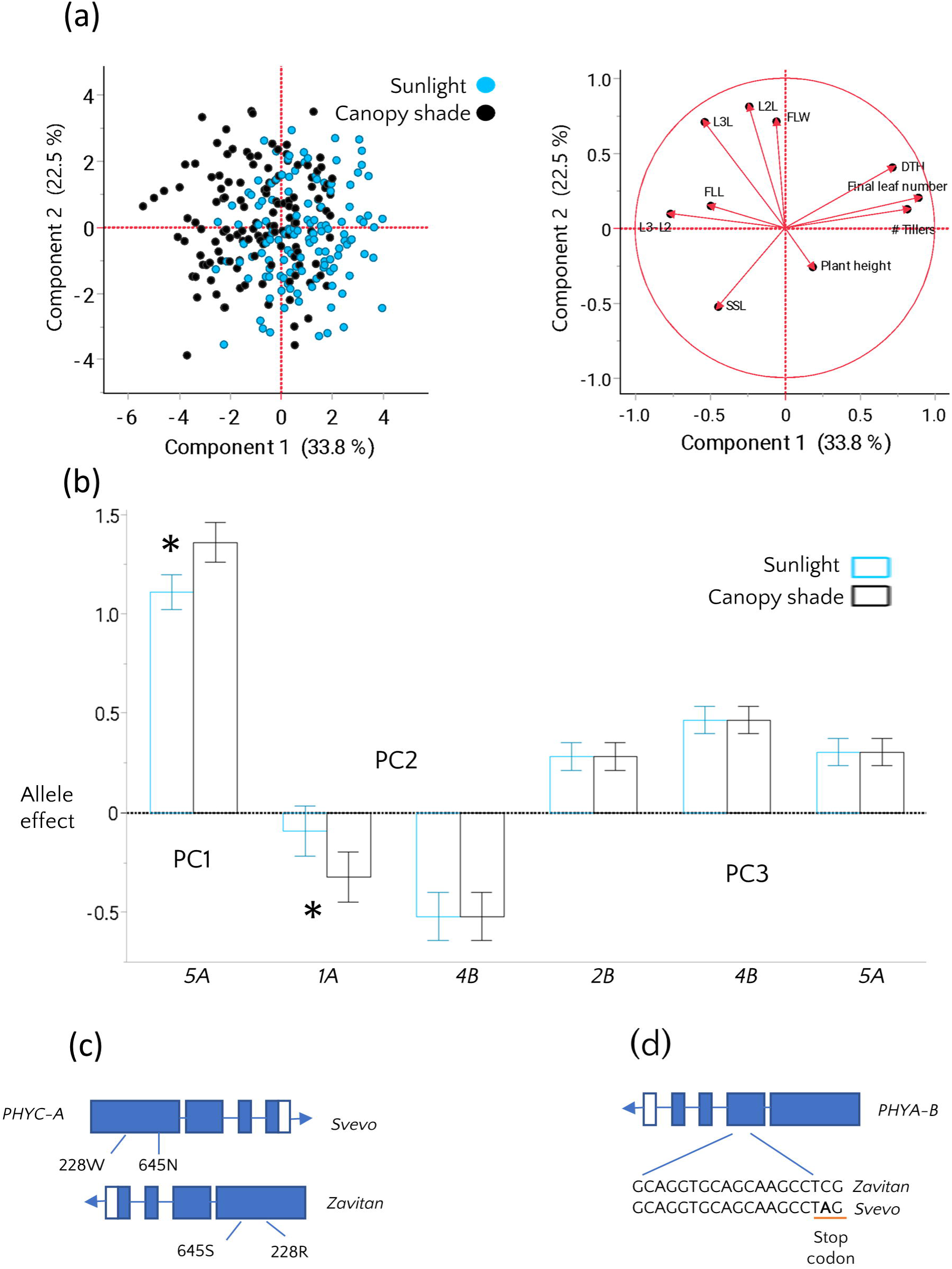
Phenotypic and genetic analysis of shade response in the SxZ RIL population. (a) principal component analysis of morphological and phenological traits under sunlight (blue) and simulated canopy shade (black). (b) Allelic effects ±SE of QTL identified for PC 1-3, demonstrating QTL-by-environment interactions between sunlight and simulated canopy shade for QTLs on chromosomes 5A and 1A. Positive allelic effects indicate increasing allele originated from Zavitan and negative values indicate increasing allele originated from Svevo. Asterisks indicate significant QTL x E interactions. (c) Schematic representation of polymorphism in PHYC-A between parental lines, including gene inversion and amino acid substitutions associated with the 5A QTL. (d) An early stop codon in PHYA-B of Svevo associated with the 4B QTL. Abbreviations: Days to heading (DTH), Specific stem length (SSL), Flag leaf length (FLL), Leaf 2 length (L2L), Leaf 3 length (L3L), Flag leaf width (FLW).

To uncover the genetic loci underlying plant architecture in our population and assess their contribution to phenotypic variation across light environments, we conducted QTL mapping of PC1–3 using linear mixed models that included environmental effects, QTL effects, and their interaction with the light environment. These interactions were explicitly tested to identify plasticity loci whose effects on phenology and architecture differed between sunlight and simulated canopy shade. PC1 phenotypes, i.e. leaf and tiller number, leaf spacing, and heading date, were associated with one major QTL on chromosome 5A, where the wild emmer allele increases tiller and leaf number, delays days to heading, and decreases the distance between leaves on the main culm. Importantly, the PC1 QTL effects were environment-dependent and were amplified under canopy shade (Fig. 1b, Table S3). Given the strong phenological loading of PC1, the observed QTL × environment interaction indicates genotype-specific differences in developmental response to canopy shade, specifically in the acceleration of development. (Fig. S4).

PC2, associated with leaf size, was regulated by two QTL on chromosomes 1A and 4B, with the elite durum alleles at both loci associated with increased leaf size. The QTL on chromosome 1A exhibited a stronger effect under shade conditions. PC3, i.e. plant height, SSL, and FLL, was influenced by three QTL: two with increasing alleles derived from the wild emmer accession and one from durum (Fig. 1b, Table S3). The QTL for PC3 on chromosomes 4B and 5A overlapped with those detected for PC1 and PC2 (Fig. 1b). These QTL did not show significant interaction with the light environment, suggesting that the observed environmental variation for PC3 is largely independent of the genotype. The 4B QTL was associated with the *Reduced-height-B1* (*Rht-B1*) locus likely reflecting known effects on plant height that contribute to variation in PC3.

### Structural variation and coding sequence polymorphisms in phytochromes are associated with plant architecture and phenology

To identify candidate genes underlying the major QTL on chromosome 5A associated with phenology and its interaction with light environment, we examined the gene content of the corresponding genomic interval. This region contains two key regulators of flowering time in wheat, *PHYTOCHROME C* (*PHYC-A*) and *VERNALIZATION-1* (*VRN-A1*). Inspection of the parental genome assemblies, supported by a PCR assay targeting one of the predicted breakpoint junctions, upstream of the *PHYC-A* coding region, revealed a previously uncharacterized ∼2 Mb inversion spanning this region, and encompassing both *PHYC-A* and *VRN-A1* (Fig. S5). The inversion was evident when examined in the improved Zavitan genome, conducted with the aid of optical maps (Zhu et al. 2019), and is further supported by a structurally similar rearrangement with comparable breakpoints in the hexaploid wheat genome of *Triticum spelta*, making it less likely to represent an assembly artifact. Such structural variation around *PHYC* is consistent with earlier reports showing that upstream regions of *PHYC* homoeologs in wheat are highly dynamic and frequently disrupted by transposable elements and other structural rearrangements (Devos et al., 2005).

Moreover, the Svevo allele of *VRN-A1* carries a ∼7.4 kb deletion in its first intron, compared to Zavitan, which has been previously shown to be associated with a dominant spring growth habit (Fu *et al*., 2005). In contrast, Zavitan carries a winter *VRN-A1* allele with an intact first intron, containing the ancestral allele in the RIP-3 region of intron 1, that strongly represses *VRN-A1* expression (Kippes et al. 2015). Furthermore, two predicted amino acid substitutions were identified in the PHYC-A protein of the wild emmer allele Zavitan compared to Svevo: one in the GAF domain (W228R) and one in the PAS-related domain (N645S, Fig. 1C), important for phytochrome nuclear import (Chen et al. 2005). Within the major-effect QTL on chromosome 4B associated with PC2 and PC3, we identified several amino acid substitutions and a premature stop codon in the *PHYTOCHROME A* (*PHYA-B*) gene of Svevo. This mutation results in a truncated PHYA-B protein lacking 241 amino acids at the C-terminus, including the histidine kinase-related domain (Fig. 1D). Interestingly, *PHYA-B* was also associated with a QTL for heading time, explaining 12% of the phenotypic variation, where RILs carrying the truncated protein from Svevo flowered on average a day later (Table S3).

To further investigate the prevalence of these polymorphisms across the tetraploid wheat gene pool, we developed PCR markers and genotyped a *Triticum turgidum* diversity panel for the identified *PHYA-B* (4B), *PHYC-A* (5A), and *VRN-A1* (5A) polymorphisms identified in the RIL population. The panel included wild emmer, domesticated emmer, modern durum, and other domesticated forms such as *T. polonicum*, *T. carthlicum, T. compositum*, and *T. aethiopicum*. The 228W allele of Svevo’s PHYC-A on 5A was rare, identified only in seven modern durum varieties (Table S4, Fig. S6). The 645S allele was present in 107 accessions, of which 94 carried the Zavitan-like inversion marker (Hap 7 in Fig. S7). Overall, we found varying frequencies of *PHYC-A* haplotypes across continents and domestication groups. The premature stop codon in *PHYA-B* on 4B was restricted to only two wild emmer accessions but was widespread among domesticated wheats. Furthermore, its frequency was higher in European genotypes (∼80%) compared to Asian genotypes (∼40%) (Table S4, Fig. S6).

Importantly, the diversity panel included accessions with the inverted/645S alleles, present both with and without the large deletion in *VRN-A1* (Fig. S7, Table S4), indicating that the structural haplotype associated with the inversion marker is not perfectly coupled with the allelic status at the *VRN-A1* locus across the diversity panel. This suggests either variation in inversion breakpoints among genotypes that affect the length of the inverted region or independent origins of *VRN-A1* intron deletions in different chromosome 5A backgrounds.

We next tested whether polymorphisms in *PHYC-A* and *PHYA-B* were associated with heading time in the tetraploid diversity panel. The 228W substitution in PHYC-A (5A) present in Svevo was found only in accessions carrying the dominant *VRN-A1* Spring allele and was associated with earlier flowering (P=0.05) compared to accessions carrying the 228R (Fig. 2a). However, we cannot rule out that variation in heading time is related to other variation in *VRN-A1,* which were not captured with the deletion marker. The inverted/645S *PHYC-A* allele was detected in both *VRN-A1* dominant and wild-type backgrounds. In the presence of the dominant *VRN-A1* spring allele, the inverted/645S allele had no detectable effect on flowering time. However, in the absence of the dominant spring allele, the inverted/645S allele was associated with delayed flowering (P<0.0001), suggesting background-dependent associations, although the effects of *PHYC-A* and linked variation at *VRN-A1* cannot be fully separated in this analysis (Fig. 2b). Moreover, the effects of the inversion and the 645S allele could not be separated (e.g., through comparison of Hap4 and Hap7) because these variants were typically co-inherited and only three Hap4 accessions had flowering time data.

**Figure 2:**
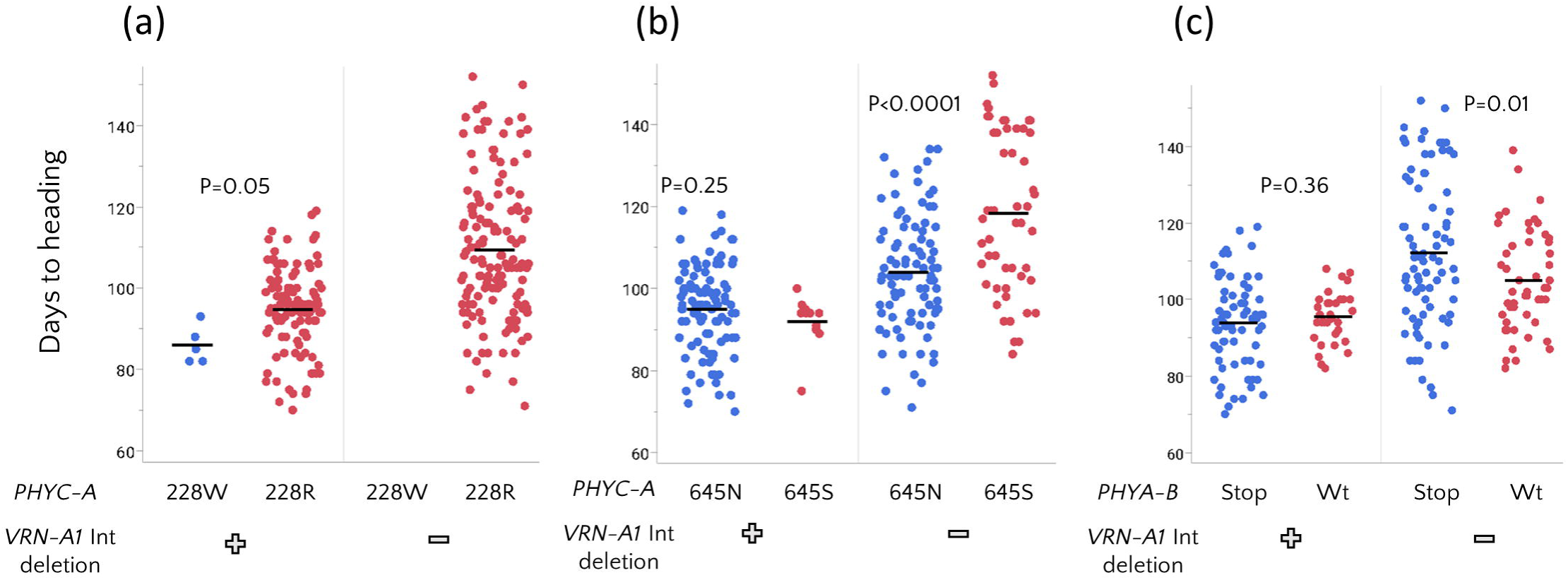
Interaction between Phy and VRN-A1 alleles affecting flowering time. (a) Earlier flowering of genotypes carrying the 228W allele. (b) Earlier flowering was associated with the 645N allele when the dominant VRN-A1 deletion was absent, but this effect was not observed when the deletion was present. (c) Early stop codon in *PHYA-B* is associated with a delay in flowering time in the absence of the dominant *VRN-A1* deletion. P Values were determined using student’s t test (a) and Welch’s t test (b-c).

A similar pattern was observed for the early stop codon in *PHYA-B* (4B): it was not associated with flowering time in accessions carrying the dominant *VRN-A1* (5A) allele, but was associated with later flowering when the dominant spring allele was absent (Fig. 2c). Overall, our findings reveal flowering time variation associated with *PHYA-B* and *PHYC-A* across different wheat genetic backgrounds, providing support for phytochrome involvement in developmental timing. However, allele frequencies varied across continents for several loci (Fig. S6), indicating geographic structuring of phytochrome and flowering-time variation that may partially confound genotype–phenotype associations.

### Natural variants in *PHYA-B* and *PHYC-A* are associated with differential expression of flowering-time genes

Based on our findings of a significant association between phytochrome genes and flowering time, we further investigated the relationship between the natural alleles of *PHYA* and *PHYC* and the expression of key flowering-time genes in the RILs. For this, we grouped the RILs by their allelic status at four flowering-time loci associated with known flowering activators (2B-*PPD-B1*, 4B–*PHYA-B*, 5A–*PHYC-A/VRN-A1*, 7B–*FT-B1*), which were identified in the current study (Tables S3, S5). This enabled us to test the effect of *PHYA-B* and 5A-*PHYC-VRN-1* while the other two loci on chromosome 2B and 7B remained fixed. Three distinct allele combinations were selected for analysis: (1) *PHYA-B^stop^*/*PHYC-A^228W^* – RILs carrying both *PHYA-B* and *PHYC-A* alleles from Svevo; (2) *PHYA-B^WT^*/*PHYC-A^228W^* – RILs carrying the *PHYA* allele from Zavitan and PHYC from Svevo; and (3) *PHYA-B^stop^/PHYC-A*^645S^ – RILs harboring the *PHYA* allele from Svevo and *PHYC* from Zavitan. Gene expression was measured in leaves at the 4^th^ leaf stage from plants grown under controlled growth chamber conditions to assess diurnal regulation associated with allelic state. qRT-PCR was conducted for *PHOTOPERIOD-1* (*PPD-1*), *VERNALIZATION-1* (*VRN-1*), and *FLOWERING LOCUS T-1* (*FT-1*) across five time points over a 24-hour period: 2, 6, 10, 18, and 22 hours (Zeitgeber Time).

Comparison between groups *PHYA-B^stop^*/*PHYC-A^228W/645N^*and *PHYA-B^WT^*/*PHYC-A^228W/645N^*, which differ in the allelic origin of *PHYA*, showed that RILs carrying the *PHYA* allele from Svevo exhibited lower expression levels of *PPD-1* and *VRN-1* across the day (6-22h) (Fig. 3a-c). These findings suggest that allelic variation at *PHYA* is associated with variation in the expression of photoperiod and vernalization pathway genes. Comparison between *PHYA-B^stop^*/*PHYC-A^228W/645N^*and *PHYA-B^stop^/PHYC-A^228R/^*^645S^ groups, which vary in their *PHYC* allele while sharing the same *PHYA* background, showed increased expression of flowering time genes across the day, the difference was evident mainly in *VRN-1* and *FT-1* in RILs carrying the *PHYC-A^228W/645N^*allele (Fig. 3d-f). Together, these results are consistent with the effect of natural variation in *PHYA* and *PHYC* on flowering time in wheat and thus provide mechanistic insight into the genetic basis of photoperiodic flowering responses in the Svevo × Zavitan RIL population.

**Figure 3:**
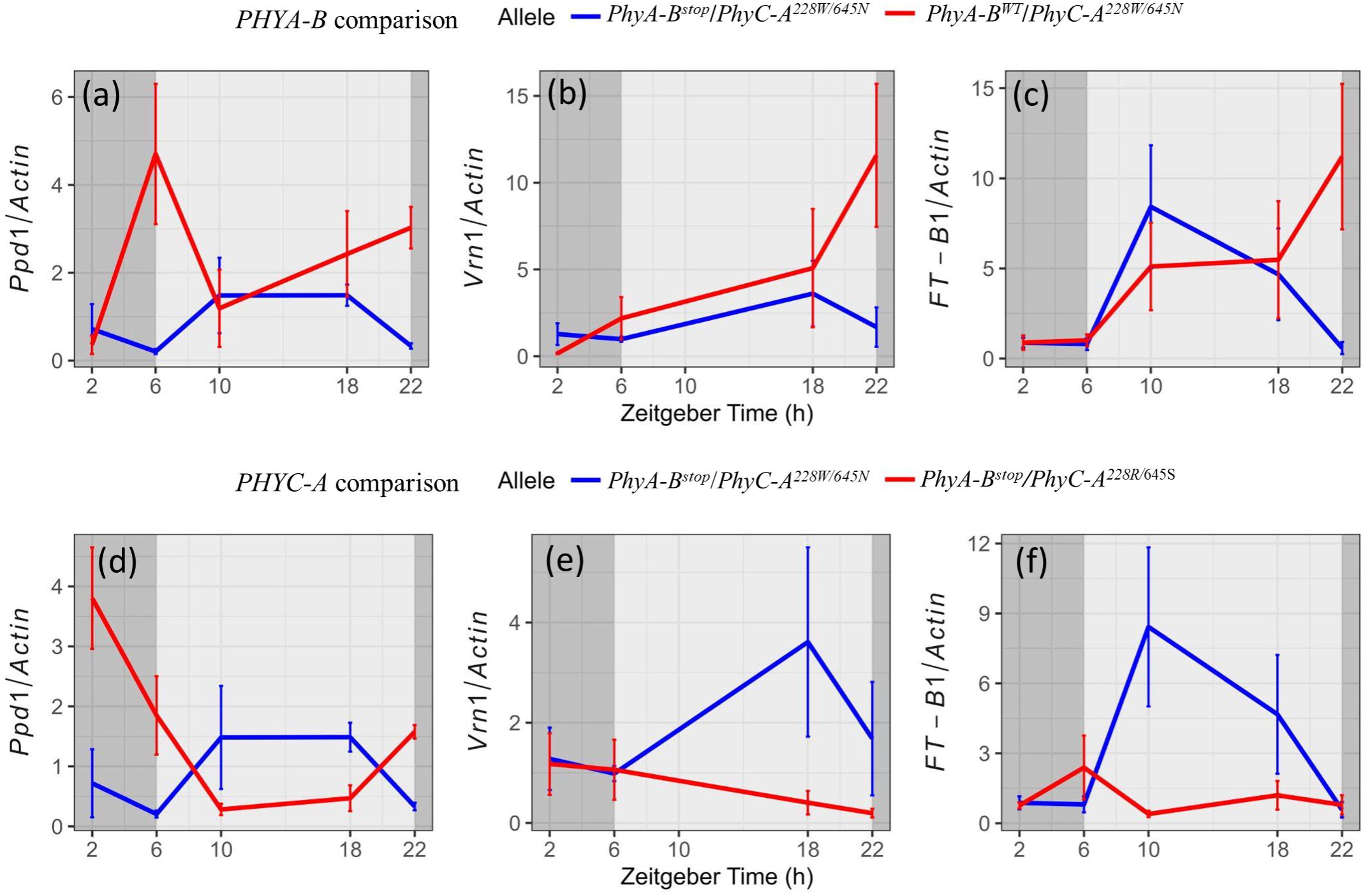
Expression of flowering time genes in RIL groups harboring contrasting PHYA-B and PHYC-A alleles. Diurnal expression profiles of flowering time genes in RILs differing in PHYA-B alleles (Svevo vs. Zavitan) while other flowering time loci fixed: (a) *PPD1*, (b) *VRN1*, (c) *FT1*. (d–f) Diurnal expression profiles of flowering genes in RILs differing in *PHYC-A* alleles (Svevo vs. Zavitan) while other flowering time loci fixed: (d) *PPD1*, (e) *VRN1*, (f) *FT1*. Each data point represent mean of 4-5 biological replicates (RILs) within each group. Error bar indicates± standard error.

### Functional characterization of *PHYA* and *PHYC* mutants reveals differences in shade-induced plasticity and plant architecture

Despite extensive evidence that deleterious mutations in *PHYC* affect flowering time, their role in modulating flowering time in response to shade remains largely unexplored. Moreover, to assess whether the premature stop codon in *PHYA-B* (4B) found in domesticated wheat may similarly affect flowering-time phenotype, we examined flowering time in TILLING-derived near-isogenic lines (NILs) carrying mutations in the background of the cultivar Kronos. We used single rather than double mutants because the extreme phenotypes exhibited by the double mutants (Chen et al., 2014; Kippes et al., 2020) are unlikely to reflect the subtle phenotypic differences observed in natural variation within a single genome. The mutants were used to assess the functional contribution of *PHYA* and *PHYC* to phenology and architecture, rather than to directly test the effects of specific natural alleles. The mutants were grown under control (sunlight) and shaded (green filter) conditions similar to those applied for the RILs. Both *PHYA-A* and *PHYA-B* mutants showed a delay in flowering time compared to Kronos, consistent with the idea that reduced *PHYA* signaling could underlie the 4B-QTL association. In the case of *PHYC*, the delay was more pronounced in the *PHYC-B* mutant (Fig. 4a), in agreement with earlier observations (Chen *et al*., 2014). Under simulated canopy shade, Kronos showed an acceleration of heading time compared to control conditions. In contrast, this acceleration was not statistically significant in PHYC mutants, supporting the idea that natural variation in PHYC may affect shade responses in addition to influencing heading date.

**Figure 4:**
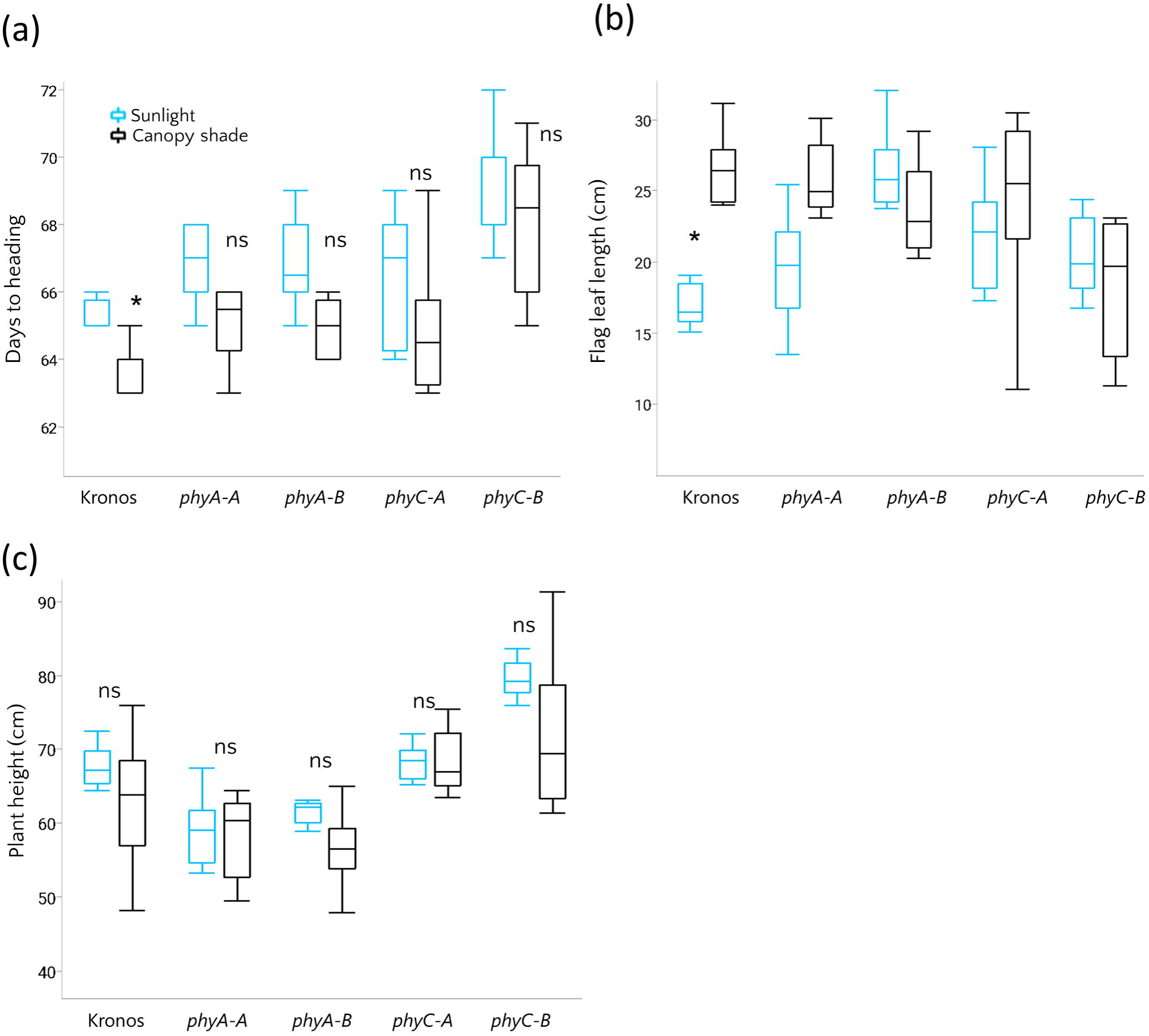
Shade response of phytochrome mutants. (a–c) Days to heading, flag leaf length, and plant height in Kronos, *PHYA-A, PHYA-B, PHYC-A*, and *PHYC-B* mutants (n=8) measured under semi-field conditions in sunlight (blue) and simulated canopy shade (black). Boxes indicate the interquartile range (25th–75th percentiles), the central line represents the median, and whiskers show the full data range. * Indicates significant differences between light treatments within each genotype, as determined by Student’s t-test.

Our QTL analysis suggested that natural variation in *PHYC* and *PHYA* is associated with variation in plant architecture. We further examined flag leaf length across genotypes and environments. Under sunlight conditions, phytochrome mutants exhibited longer flag leaves than the wild type (Fig. 4b). However, under simulated canopy shade, flag leaf elongation was observed only in the *PHYA-A* mutant and in Kronos, while the other phytochrome mutants showed no differences compared to sunlight conditions (Fig 4b). This indicates that phytochromes regulate both leaf size and its plastic response to shade in wheat, and their absence leads to an elongated phenotype largely independent of light cues. Both *PHYA-A* and *PHYA-B* (chr 4) mutants exhibited a significant reduction in plant height (at maturity) relative to Kronos, suggesting that *PHYA* promotes vertical growth. In contrast, *PHYC* mutants were taller (Fig. 4c), indicating opposing effects of different phytochromes on plant height.

## Discussion

To improve crop adaptation in dense stands, it is vital to understand how plants integrate light signals with their developmental programs. In this study, we identified naturally occurring genetic variants in the phytochrome genes PHYA and PHYC associated with notable differences in wheat phenology, architecture, and plastic responses to shade. Through a combination of QTL mapping, structural variant analysis, expression profiling, and functional analysis using TILLING mutants, we show that genetic variation in phytochromes is associated with shade-responsive morphological adaptations and developmental timing in the tetraploid wheat gene pool, consistent with a key role in these processes.

### Natural variation at the 5A Locus: The role of *PHYC* and structural inversions

The 5A locus, which is associated with shade responses, contains two major regulators of flowering time, *PHYC* and *VRN-1*. The *VRN-A1*-dependent effect of *PHYC* observed in our study aligns with the genetic models of flowering time regulation, in which *PHYC* acts upstream of *VRN-1* as a light sensor and is required for proper activation of the photoperiod pathway (Chen *et al*., 2014). Here, the unique *PHYC-A* haplotype in Svevo may confer enhanced sensitivity to altered light quality and may contribute to the genotype-specific, shade-associated shifts in flowering time (Fig. S4). These findings are consistent with a recent study demonstrating that barley *PHYC* function is highly context-dependent, with its effects on flowering time varying across environments and genetic backgrounds through epistatic interactions with key flowering genes (Okuma et al. 2026).

At the protein level, the positively charged arginine (R) at position 228, which is conserved among PHYC proteins across plant species, is replaced by tryptophan (W), a bulky, hydrophobic aromatic residue. Given the magnitude of this physicochemical change, it could possibly affect PHYC structure or signaling, but this interpretation requires targeted functional assays. The affected GAF domain provides the binding pocket for the bilin chromophore that is essential for triggering the conformational changes between the active and inactive phytochrome states, and amino acid changes in this region have been suggested to enhance phytochrome activity and flowering responses (Nishida *et al*., 2013; Pankin *et al*., 2014).

In addition to coding variation in *PHYC-A*, the 5A locus is characterized by a large structural rearrangement consistent with an inversion between Svevo and Zavitan, encompassing both *PHYC-A* and *VRN-A1*, which may suppress recombination in this interval. We inferred this inversion from comparison of genome assemblies of Svevo and Zavitan which was further validated using a PCR assay that targets the inversion break point. However, in the diversity panel, the structural state was inferred using the breakpoint-associated marker, and we do not know whether all accessions share identical inversion breakpoints or lengths. Moreover, the presence of Zavitan-like structural haplotypes with and without the *VRN-A1* intron deletion indicates that linkage between these loci is incomplete across the diversity panel, suggesting either variation in inversion breakpoints or independent origins of *VRN-A1* alleles in different genetic backgrounds. Previous studies of the upstream regions of wheat *PHYC* homoeologs have revealed frequent insertions of transposable elements and substantial structural divergence among genomes, suggesting that different genomic regions have distinct evolutionary trajectories contributing to a mosaic genome structure (Devos et al., 2005). This may help explain the complex patterns of linkage and allele distribution observed at the *PHYC-A–VRN-A1* locus in our study.

Such structural variants can facilitate the maintenance of co-adapted allele combinations by preventing the breakup of favorable haplotypes (Lowry and Willis, 2010; Todesco *et al*., 2020; Wellenreuther et al. 2025). This specific inversion of wheat may preserve specific allele combinations for photoperiod and vernalization, and might have played an adaptive role by creating specific haplotypes that ties together light sensing and vernalization requirements with shade-induced plasticity. The intermediate frequency, broad phylogenetic distribution, and different frequencies across continents (Table S4, Fig. S6) suggest that this structural variation represents standing variation that predates domestication and has been maintained across evolutionary and breeding history, perhaps reflecting an environment-dependent haplotype selection.

### PHYA-B as a candidate contributor to flowering time variation

The premature stop codon identified in *PHYA-B* is predicted to remove a substantial portion of the C-terminal region, including the histidine kinase-related domain required for complete signaling functionality (Krall and Reed, 2000). Interestingly, this allele is rare in wild emmer but common in domesticated wheat, indicating it may have increased in frequency during domestication or improvement (via selection or genetic drift). Moreover, the premature stop codon showed a significant association with geographic distribution and was more frequent in European genotypes compared to Asian and African genotypes, suggesting that it may be linked to environmental adaptation. In our RIL population, natural variation at the *PHYA* locus on chromosome 4B was associated with variation in heading time and plant height. However, this gene was genetically linked to the *Reduced height-B1* (*Rht-B1*) locus, which encodes a DELLA protein that suppresses GA-mediated growth. In the RIL population, *Rht-B1* accounts for a large proportion of the variation in plant height (Golan *et al*., 2024) and is therefore likely to mask any detectable effects of *PHYA* on this trait, making it difficult to resolve independent contributions of *PHYA-B* to plant height variation.

By comparison, multiple studies have reported flowering-time QTL co-localizing with the *Rht-B1* region (Peleg *et al*., 2009; Sanna *et al*., 2014; Milner *et al*., 2016; Giunta *et al*., 2018; Gupta *et al*., 2020). However, Milner et al., (2016) did not detect a plant height QTL at this locus, and the study by Peleg et al., (2009) was conducted using two tall parental lines lacking major dwarfing alleles. Moreover, *Rht-B1* has little or no direct effect on flowering time (Langer *et al*., 2014). In contrast, several studies show that *PHYA* mediates developmental and flowering time responses to low light and far-red light enrichment in *Arabidopsis* (Quail, 2002; Franklin and Quail, 2010; Seaton *et al*., 2018), Medicago (Jaudal *et al*., 2020), soybean (Lin *et al*., 2022) and rice (Takano *et al*., 2001, 2005*b*,*a*). Taken together, these observations make *PHYA* a strong candidate contributor to the flowering-time QTL detected in this region, while the strong effect of *Rht-B1* on plant height likely obscures potential plant height-related effects of *PHYA*.

### Phytochromes as diurnal regulators of the flowering pathway

Further, expression analysis across multiple time points revealed that allelic differences at *PHYA* and *PHYC* are associated with the transcriptional dynamics of key flowering genes, including *PPD-1, VRN-1*, and *FT-1*. RILs containing the Svevo *PHYA-B^stop^* allele, which encodes a putative truncated protein, displayed reduced *PPD-1* and *VRN-1* expression, particularly at the end of the day. However, further validation linking natural variation in *PHYA* to flowering time and mechanistic models incorporating *PHYA* in wheat flowering time pathways are still lacking.

Allelic variation at *PHYC* was associated with the daytime expression of *VRN-1* and *FT-1*, indicating a significant role for *PHYC* in regulating light-dependent activation of subsequent flowering pathways. These results are consistent with the previous models of flowering time in cereals, where *PHYB* and *PHYC* regulate flowering time by modulating the diurnal expression of *PPD-1*, which in turn activates *FT-1* and *VRN-1* (Chen *et al*., 2014). Together, these data link allelic state at *PHYA*/*PHYC* to diurnal differences in expression of core flowering regulators.

### Functional validation and implications for wheat improvement

TILLING mutants confirmed that both *PHYA* and *PHYC* can influence wheat growth and development. The delay in heading observed in *PHYA* and *PHYC* mutants in the Kronos background lacking the large intron deletion in *VRN-A1* supports the allelic associations observed in the diversity panel and underscores the requirement of both phytochromes for proper induction of flowering pathways under long days. The stronger phenotype of the *PHYC-B* mutant, compared with *PHYC-A*, likely reflects both functional differences between the homoeologs and the possibility of allele-specific effects in the TILLING lines, and is consistent with prior reports identifying *PHYC-B* as the dominant phytochrome regulating *PPD-1* and *FT-1* expression in cv. Kronos (Chen *et al*., 2014). Moreover, *PHYA* and *PHYC* mutants displayed altered vegetative architecture, including plant height and flag leaf length. These findings align with previous studies showing that phytochromes regulate internode elongation and leaf expansion through modulation of auxin and GA-related shade-avoidance pathways (Pierik and De Wit, 2014; Li *et al*., 2024, Tao et al. 2008).

Together, our findings reveal that natural alleles of *PHYA* and *PHYC* are associated with phenotypic plasticity and flowering-time variation across the tetraploid wheat gene pool. Shade avoidance traits can influence early vigor, canopy structure, weed competitiveness, and performance in dense plantings, traits relevant to modern agriculture. The identification of new alleles associated with developmental timing and plasticity provides opportunities for refining flowering time under diverse environments or altering plant architecture to improve resource-use efficiency. Importantly, the widespread presence of the truncated *PHYA* allele in domesticated wheats strongly suggests that altered photoreceptor function may have contributed to wheat diversification and adaptation to new environments. The structural haplotypes involving *PHYC-A* and *VRN-A1* represents a naturally occurring variant identified in the parental genomes and inferred using a marker assay across the tetraploid wheat gene pool. It occurs in both wild and domesticated accessions, and its effects on flowering, development, and adaptation require further ecological and agronomic investigation. Overall, our study connects previously unknown variation in wheat phytochromes with plant development and shade avoidance.

## Acknowledgements

The authors thank Jorge Dubcovsky for providing grains of the backcrossed TILLING mutants and for critical review of an earlier version of the manuscript. We also thank Assaf Distelfeld and Zvi Peleg for providing the RIL population. We are grateful to Yongyu Huang for support with genomic analyses and visualization of the inversion in Svevo and Zavitan. We further thank Enk Geyer and his team for their support during the growing season, Corinna Trautewig, Kerstin Wolf, and Ellen Weiss for assistance with plant management and processing, and all the members of the plant architecture group at IPK for fruitful discussions. Infrastructure and financial support for this project were provided by the IPK core budget as part of a seed-funding project to establish joint scientific links between CEPLAS (Cluster of Excellence on Plant Sciences) and IPK. While conducting parts of this study GG received financial support from a postdoctoral fellowship of the Alexander-von-Humboldt Foundation (AvH) and TS received funding from the German Research Foundation (DFG), grant no. SCHN 768/20-1.

## Supplementary figures

**Figure S1: Correlation matrix of phenotypic traits. Heatmap showing pairwise Pearson correlation coefficients.** (a) Corresponding significance levels (P-values) are shown in the right matrix. (b) Color intensity reflects the magnitude and direction of correlations, with darker shades indicating stronger positive or negative associations.

**Figure S2: Partial contributions of the trait phenotypes recorded under sunlight and simulated canopy shade to principal components 1-3.**

**Figure S3: Phenotypic variation under sunlight and shade.** PC scores (PC1–PC3) derived from multivariate analysis of phenotypic traits plotted for plants grown under sunlight (blue) versus simulated canopy shade (black). Each point represents an individual RIL. Asterisks indicate significant difference (P<0.05) between treatments following analysis of variance. Mean diamonds indicate the 95% confidence intervals for the treatment means; the center line represents the mean, and the vertical extent denotes the confidence interval.

**Figure S4: Genotype by environment interaction of heading time**. Mean days to heading for RILs carrying the different 5A QTL alleles when grown under natural sunlight and simulated canopy shade. Error bars indicate ± SE.

**Figure S5. Identification of the inversion on chromosome 5A.** (a) Alignment of the 5A pseudomolecules of Svevo and Zavitan. (b) A PCR marker (Table S1) targeting the proximal inversion break point was used to screen for the structural rearrangement in the tetraploid diversity panel that represents Svevo (S) and Zavitan (Z).

**Figure S6. Geographic and domestication-associated variation in phytochrome and VRN-A1alleles in tetraploid wheat.** Allele frequencies are shown as proportions within each group. (a–e) Distribution of alleles across continents for (a) structural haplotype,(b) PHYC-A at position 228 (R/W), (c) PHYC-A at position 645 (N/S), (d) VRN-A1 intron 1 deletion, and (e) PHYA-B allele (stop vs wild type). (f–j) Corresponding allele frequencies grouped by domestication status (domesticated vs wild emmer accessions) for the same loci.

**Figure S7: Haplotype classes on chromosome 5A defined by polymorphic sites in *PHYC-A* and *VRN-A1* of Svevo and Zavitan**. Haplotypes are defined based on combinations of *PHYC-A* coding variants and *VRN-A1* intron deletion status. Structural configuration (Svevo-like vs Zavitan-like) was inferred using marker assays targeting breakpoint regions (Fig. S5). Gene orientation is shown based on reference genome assemblies of Svevo and Zavitan, and for other genotypes, represents inferred structural configuration.

## Supplementary Tables

**Table S1:** Details of the primers used for genotyping and qPCR with their PCR protocols in this study.

**Table S2:** Mixed linear models for morphological and phenology traits examined under sunlight and canopy shade.

**Table S3:** QTL analysis data.

**Table S4:** Genotyping results for *PHYC-A, VRN-A1* and *PHYA* polymorphisms: each accession in the diversity panel was assigned the corresponding parental allele identified in the RIL (SxZ) population.

**Table S5:** Genotypic characterization of four flowering time loci across three genetic groups of Recombinant Inbred Lines used for gene expression analysis.

