## Supplementary figures for "Structural and coding variation in *PHYTOCHROMES A* and *C* underlies differences in flowering time and shade avoidance in wheat"

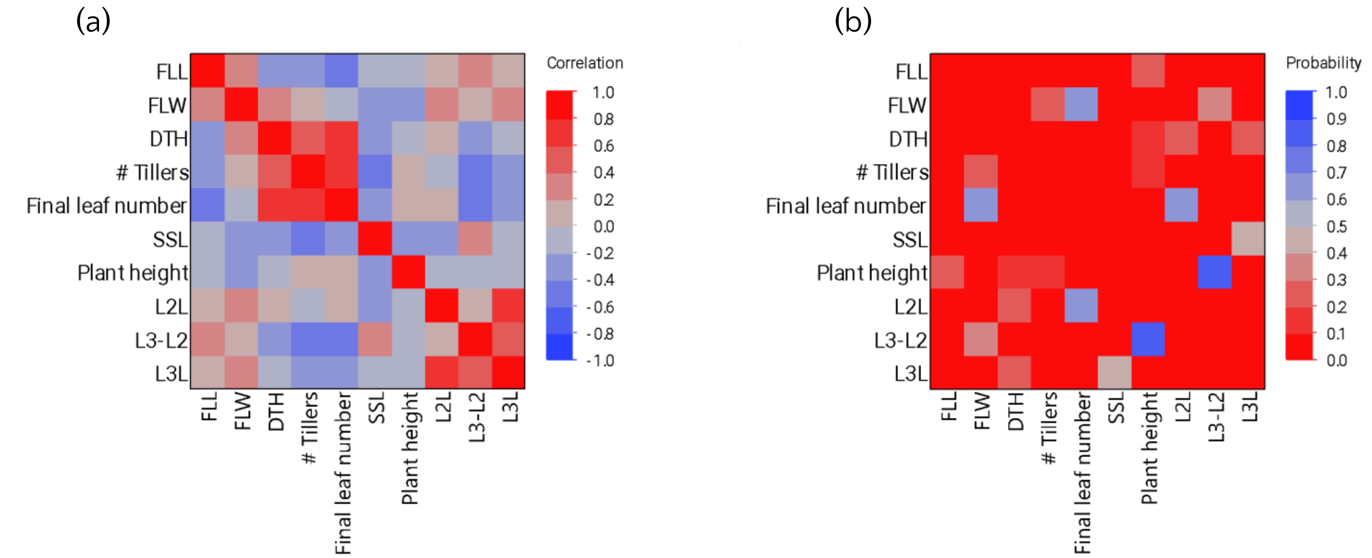


**Fig. S1 Correlation matrix of phenotypic traits**. Heatmap showing pairwise Pearson correlation coefficients (a). Corresponding significance levels (P-values) are shown in the right matrix (b). Color intensity reflects the magnitude and direction of correlations, with darker shades indicating stronger positive or negative associations. Abbreviations: Days to heading (DTH), Specific stem length (SSL), Flag leaf length (FLL), Leaf 2 length (L2L), Leaf 3 length (L3L), Flah leaf width (FLW).

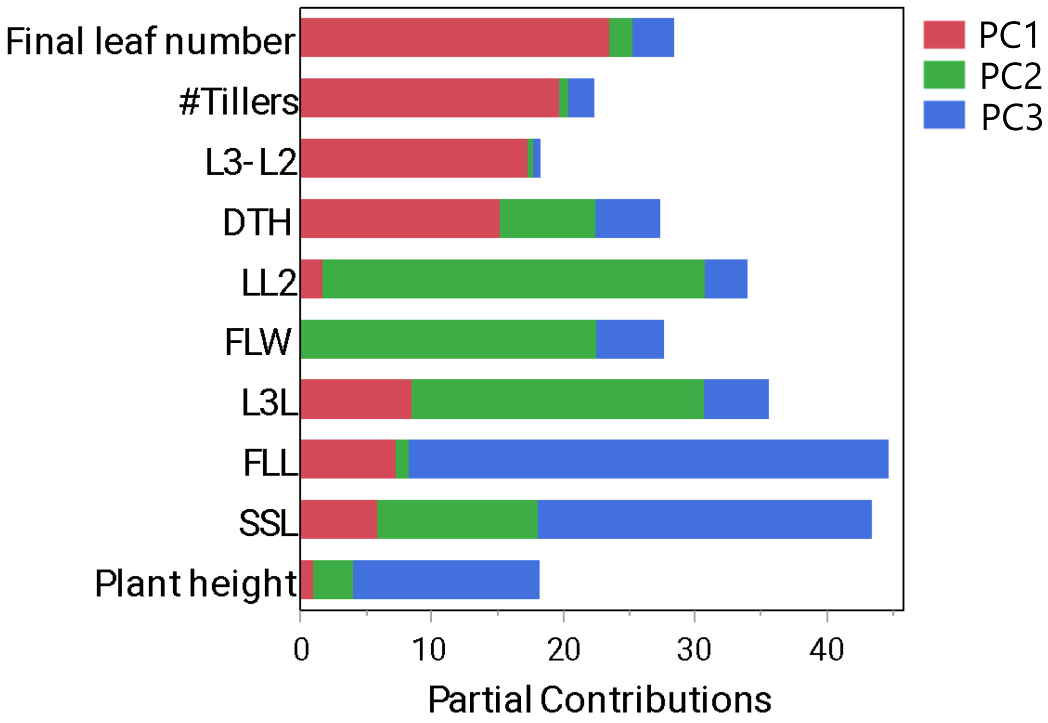


**Fig. S2 Partial contributions of the trait phenotypes recorded under sunlight and simulated canopy shade to principal components 1-3.** Abbreviations: Days to heading (DTH), Specific stem length (SSL), Flag leaf length (FLL), Leaf 2 length (LL2), Leaf 3 length (L3L), Flah leaf width (FLW).

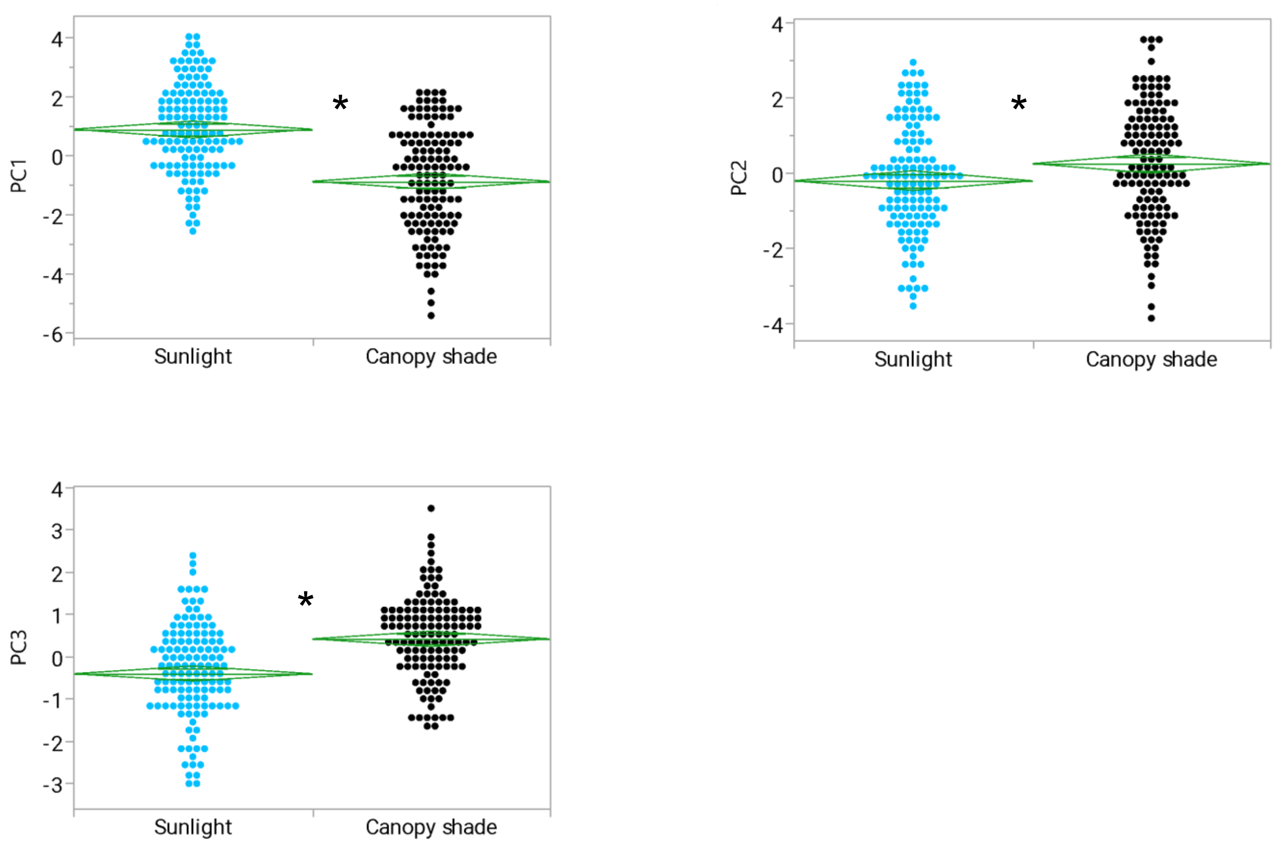


**Figure S3. Phenotypic variation under sunlight and shade**. PC scores (PC1–PC3) derived from multivariate analysis of phenotypic traits plotted for plants grown under sunlight (blue) versus simulated canopy shade (black). Each point represents an individual RIL. Asterisks indicate significant difference ($P<0.05$) between treatments following analysis of variance. Mean diamonds indicate the 95% confidence intervals for the treatment means; the center line represents the mean, and the vertical extent denotes the confidence interval.

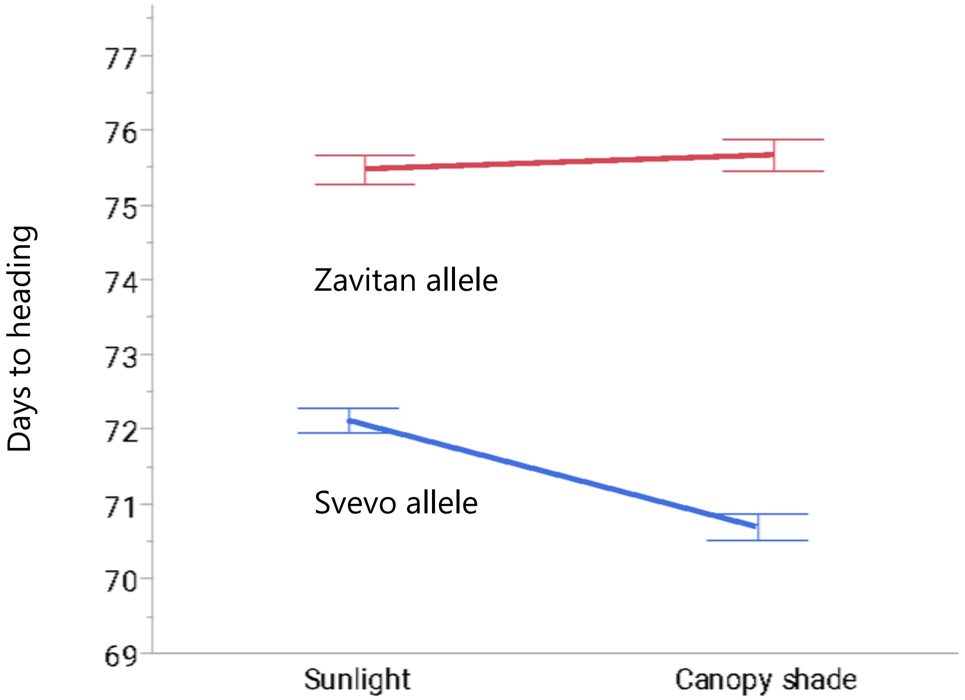


**Figure S4. Genotype by environment interaction of heading time**. Mean days to heading for RILs carrying the different 5A QTL alleles when grown under natural sunlight and simulated canopy shade. Error bars indicate ± SE.

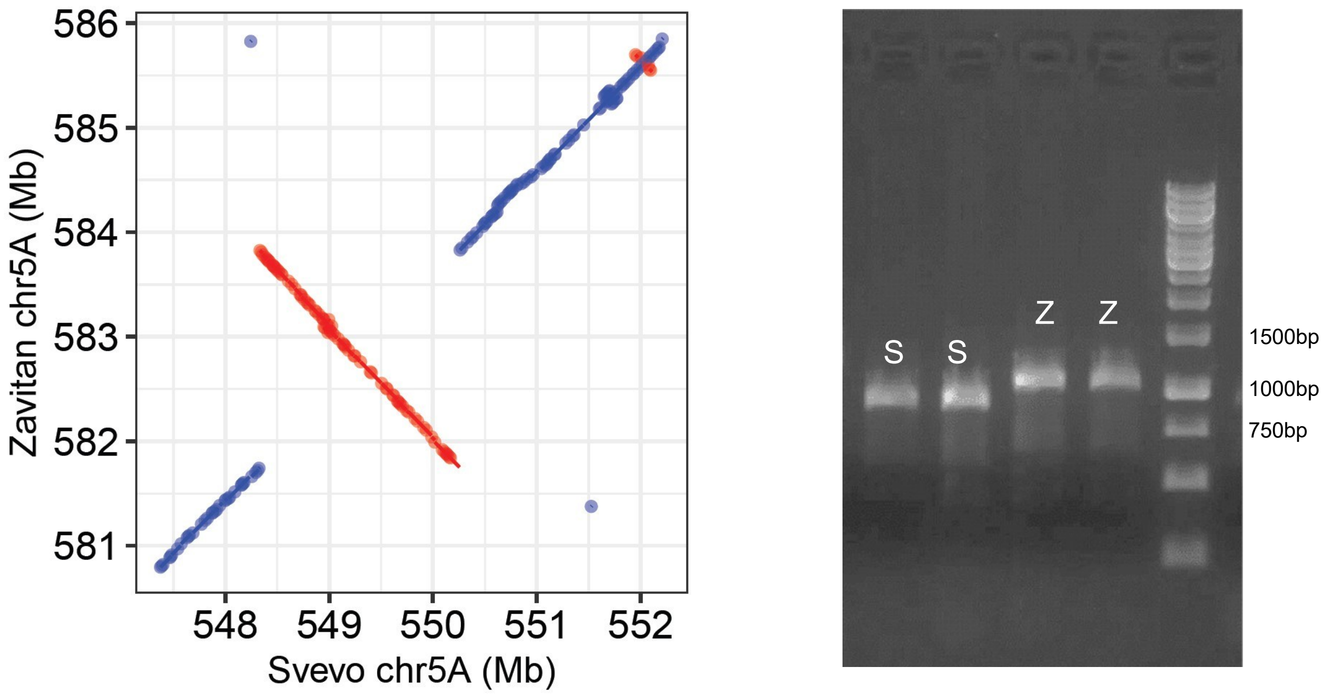


**Figure S5. Identification of the inversion on chromosome 5A.** (a) Alignment of the 5A pseudomolecules of Svevo and Zavitan. (b) A PCR marker (Table S1) targeting the proximal inversion break point was used to screen for the structural rearrangement in the tetraploid diversity panel that represents Svevo (S) and Zavitan (Z).

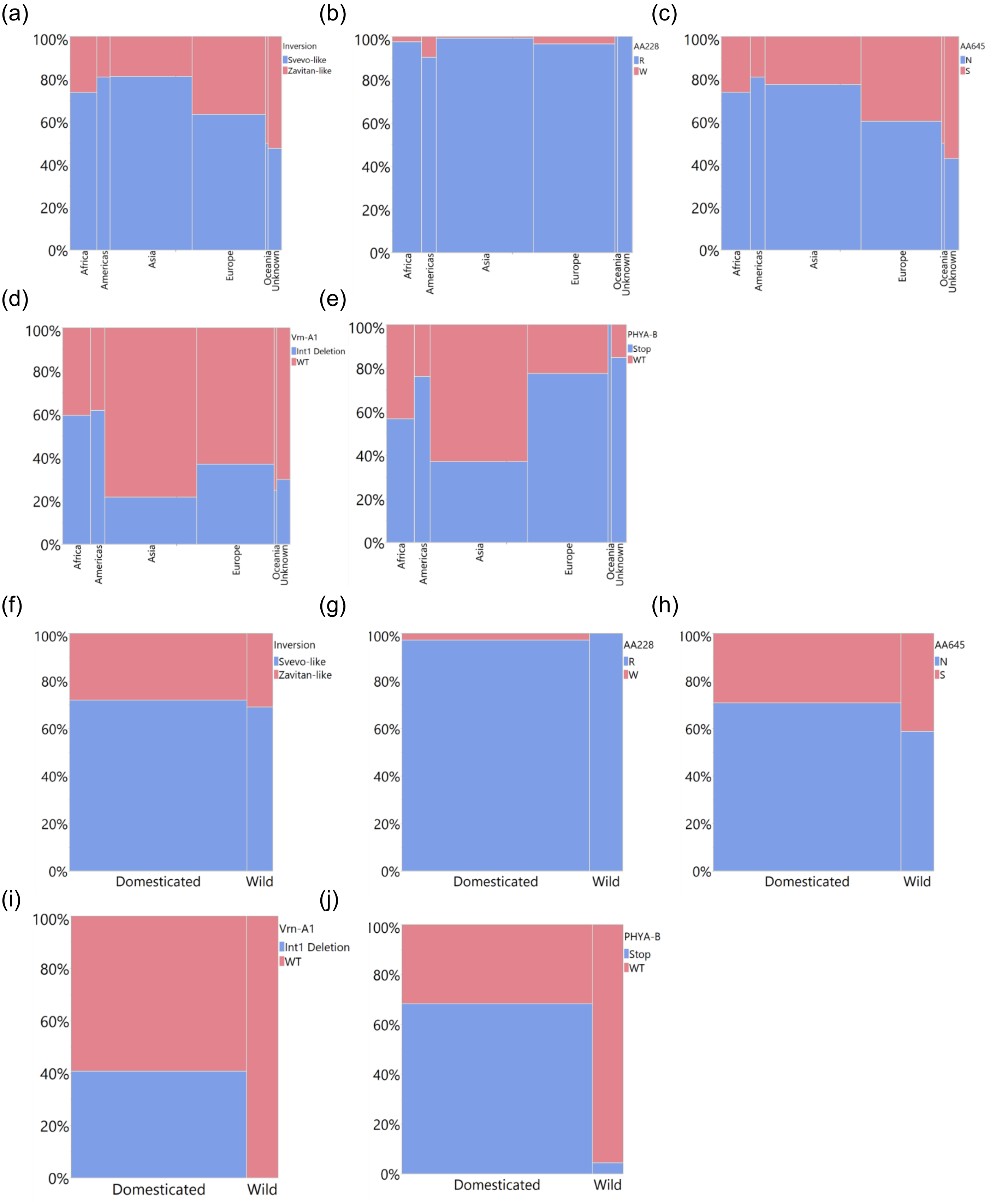


**Figure S6. Geographic and domestication-associated variation in phytochrome and *VRN-A1*alleles in tetraploid wheat**. Allele frequencies are shown as proportions within each group. (a–e) Distribution of alleles across continents for (a) structural haplotype,(b) PHYC-A at position 228 (R/W), (c) PHYC-A at position 645 (N/S), (d) *VRN-A1* intron 1 deletion, and (e) PHYA-B allele (stop vs wild type). (f–j) Corresponding allele frequencies grouped by domestication status (domesticated vs wild emmer accessions) for the same loci.

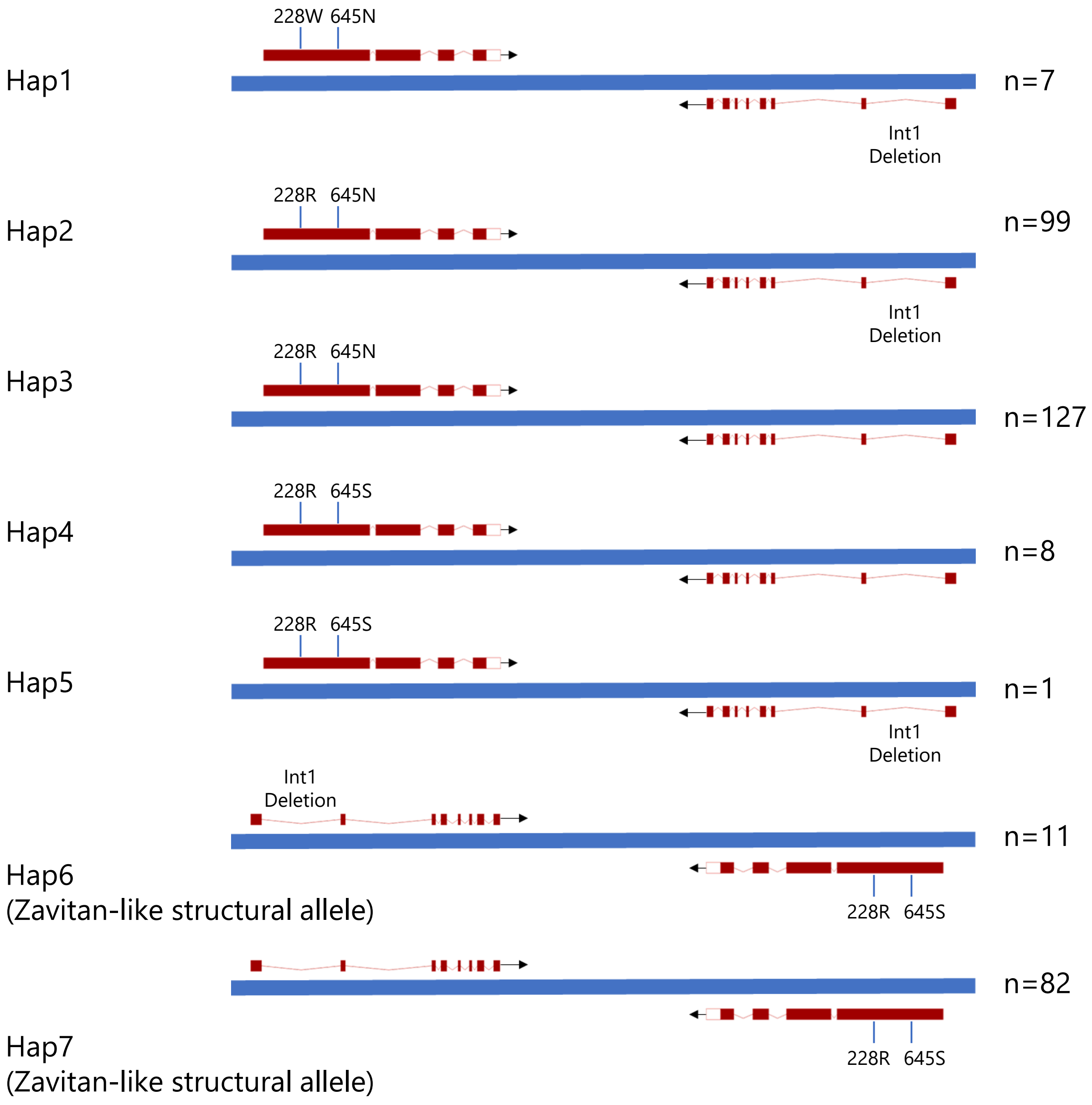


**Figure S7. Haplotype classes on chromosome 5A defined by polymorphic sites in *PHYC-A* and *VRN-A1* of Svevo and Zavitan.** Haplotypes are defined based on combinations of *PHYC-A* coding variants and *VRN-A1* intron deletion status. Structural configuration (Svevo-like vs Zavitan-like) was inferred using marker assays targeting breakpoint regions (Fig. S5). Gene orientation is shown based on reference genome assemblies of Svevo and Zavitan, and for other genotypes, represents inferred structural configuration.
